# Alcohol dependence modifies brain networks activated during abstinence and reaccess: a c-fos-based analysis in mice

**DOI:** 10.1101/2022.08.26.505400

**Authors:** Alison V. Roland, Cesar A.O. Coelho, Harold L. Haun, Carol A. Gianessi, Marcelo F. Lopez, Shannon D’Ambrosio, Samantha N. Machinski, Christopher D. Kroenke, Paul W. Frankland, Howard C. Becker, Thomas L. Kash

## Abstract

High-level alcohol consumption causes neuroplastic changes in the brain that lead to negative affective and somatic symptoms when alcohol is withdrawn, promoting relapse drinking. We have some understanding of these plastic changes in defined brain circuits and cell types, but unbiased approaches are needed to explore broader patterns of adaptations. Here, we employed whole-brain c-fos mapping and network analysis to assess how brain-wide patterns of neuronal activity are altered during acute alcohol abstinence and reaccess in a well-characterized model of alcohol dependence. Mice underwent four cycles of chronic intermittent ethanol vapor exposure (CIE) with alternating weeks of voluntary alcohol drinking, and a subset of mice underwent forced swim stress (FSS) prior to drinking sessions to further escalate alcohol consumption. After four CIE cycles, brains were collected from mice in each group either 24 hours (abstinence) or immediately following a one-hour period of alcohol reaccess. Brains from CIE mice during acute abstinence displayed widespread neuronal activation relative to those from AIR mice, independent of FSS, and this increase in c-fos was reversed by reaccess drinking. For network analysis, mice were then classified as high or low drinkers (HD or LD). We computed Pearson correlations for all pairs of brain regions and used graph theoretical methods to identify changes in network properties associated with high-drinking behavior. Network modularity, a measure of network segregation into communities, was increased in HD mice after alcohol reaccess relative to abstinence. Within-community strength and diversity measures were computed for each region and condition, and highly coactive regions were identified. One high-diversity region, the cortical amygdala (COA), was further interrogated using a chemogenetic approach. COA silencing in CIE mice reduced voluntary drinking, validating our network analysis and indicating that this region may play an important but underappreciated role in alcohol dependence.

## Introduction

Alcohol use disorders (AUD) are among the most common mental health conditions in developed countries. The transition from casual alcohol use to dependence results from long-term high-level alcohol consumption, which promotes a host of persistent molecular and cellular neuroadaptations across the brain (1,2). While initial alcohol use is motivated by its rewarding properties, pathological drinking behavior is thought to be driven in part by a constellation of negative affective and somatic symptoms that arise when alcohol is withdrawn. Alcohol-induced plasticity of central stress systems is fundamental to this transition, and altered stress responses may be a predisposing factor in relapse drinking (1,2).

Rodent models of high-level alcohol exposure have enabled mechanistic studies of this plasticity (3). One such model is the chronic intermittent ethanol (CIE) vapor paradigm, which provides a robust method of inducing alcohol dependence in rodents. Although differences have been reported based on strain and duration of exposure, mice exposed to CIE generally display numerous withdrawal-like symptoms, including enhanced seizure susceptibility (4), locomotor changes (5), affective disturbances (6), and increased voluntary alcohol drinking (7,8). CIE-induced increases in drinking can be reliably escalated by administering forced swim stress (FSS) four hours prior to drinking sessions (9–11), and this CIE/FSS model has greatly facilitated the study of stress-alcohol interactions in mice.

While these and other models have generated important insight into key neurochemicals and circuits that contribute to alcohol dependence, existing studies are biased toward brain regions already established to play a role in addiction (12), leaving a wide knowledge gap concerning other potentially influential regions. The recent implementation of unbiased whole-brain approaches and graph theoretical methods has begun to bridge this gap (13). Network-based methods can provide insight into how the brain produces complex cognitive states and behaviors, which arise from coordinated activity among groups or “communities” of brain regions (14). A recent study employed single-cell whole-brain imaging and graph theory to assess the functional brain networks associated with abstinence from alcohol in CIE mice (15). Relative to casual drinkers and alcohol-naïve mice, CIE mice displayed a decrease in network modularity after one week of abstinence, indicating that alcohol-induced plasticity is reflected in broader network-level changes.

In the present study, we further explored the outcomes of high-level alcohol exposure on brain-wide c-fos expression and functional brain networks, implementing a distinct methodological approach to construct our networks. We explored this in the well-characterized CIE/FSS model of alcohol dependence to gain insight into how stress modulates brain-wide activity patterns to escalate drinking. C-fos expression was examined at two time points: acute abstinence (24h), and immediately following a 1h alcohol reaccess period; the latter allowed us to assess how dependence alters the effects of acute alcohol consumption on neural activity. Whole-brain immunostaining for c-fos was performed using iDISCO, and c-fos-positive cell counts for 110 brain regions were obtained using Clearmap automated analysis. Interregional Pearson correlations were computed to identify functional networks based on patterns of coactivation among brain regions. We then constructed the functional networks representing the neural state during abstinence and reaccess in groups selected for low-drinking (LD) and high-drinking (HD) behavior. We compared global network properties, and for each brain region we also captured two community-based properties, within-community strength (wcs) and diversity (h), to understand how individual regions change their connectivity after alcohol exposure. While wcs captures the strength of a region’s connectivity within its assigned community, h captures the strength of its connectivity to regions of other communities. We identified several high-diversity brain regions in abstinent HD mice whose diversity coefficient was normalized by reaccess drinking. Due to their strong connectivity to multiple brain regions, these regions are thought to exert a disproportionate influence on network activity (14). To test the hypothesis that the activity of these regions influences drinking behavior in CIE mice during acute abstinence, we then used a chemogenetic approach to interrogate one of these targets, the cortical amygdala.

## Materials and Methods

### Subjects

For CIE/FSS studies, male C57BL/6J mice (10 weeks of age) were obtained from Jackson Laboratories (Bar Harbor, ME). For chemogenetic experiments, 7-week-old male C57BL/6J mice were obtained in-house from the University of North Carolina. All mice were individually housed, maintained on a 12-h reverse light/dark cycle, and provided free access to food and water throughout the duration of the experiments. CIE/FSS and brain tissue harvest protocols were approved by the Medical University of South Carolina Institutional Animal Care and Use Committee. Follow-up CIE and chemogenetic studies were approved by the Institutional Animal Care and Use Committee of the University of North Carolina. All experiments were consistent with guidelines of the NIH Guide for the Care and Use of Laboratory Animals.

### Chronic intermittent ethanol exposure

During a 6-week baseline drinking period, mice had access to a bottle containing alcohol (15% v/v) for 1 h daily (M-F) beginning 4h into the dark cycle. Mice were then divided into two groups based on the average intake in the final baseline week. Mice were exposed to either room air (AIR) or alcohol vapor (CIE) for four consecutive days (16 h on/8 h off) on alternating weeks, with interleaved test drinking weeks. Blood was collected from the retroorbital sinus or tail once during each vapor week for determination of blood ethanol content (BEC) using an Analox AM1 Analyser (Analox Instruments, UK). During test drinking weeks, mice had daily (M-F) 1h access to alcohol to assess voluntary consumption.

### Forced swim stress

During intermittent voluntary drinking weeks, subgroups of AIR and CIE mice underwent forced swim stress four hours prior to each drinking session as previously described (9,11). Briefly, mice were placed into a glass cylinder containing 23–25° C water at a depth of 20 cm for 10 minutes. Mice were placed back into their home cage, which was situated on a heated pad, for 5 minutes, and then returned to the colony room. Glass cylinders were cleaned between subjects and nonstressed mice remained in their home cages for the duration.

### Brain Tissue Collection

Following the fourth alcohol vapor cycle, mice were provided 1h access to alcohol on days 1 and 2 of test 4. On day 3, FSS groups underwent forced swim stress, and 5h later a subgroup of mice from all treatment groups were deeply anesthetized with urethane (1.5 mg/kg i.p.) and transcardially perfused with 10 mL PBS followed by 10 mL 4% paraformaldehyde (PFA) in phosphate buffered saline (PBS). The remaining mice were provided 1h access to alcohol and perfused 1h after termination of the drinking session. Brains were immediately extracted following perfusion, post-fixed for 24 h at 4C in 4% PFA, and transferred to PBS for long-term storage prior to immunostaining.

### iDISCO whole-brain immunostaining

Brains were hemisected approximately 1 mm lateral to the midline, and left and right hemispheres were collected alternately. Whole-brain immunostaining for c-fos was performed according to published methods (16). Briefly, hemisected brains were dehydrated in a series of 20%, 40%, 60%, 80%, and 100% methanol in ddH_2_O (1h each). Brains were bleached overnight in 5% hydrogen peroxide in methanol, then rehydrated in a reverse series of 100-20% methanol in ddH_2_O. Following three washes in PBS, brains were washed in PBS/0.2%Triton-X-100, permeabilized for 2 days at 37C in a solution containing 0.3M glycine and 20% DMSO in PBS/0.2% Triton-X-100, blocked for 2 days at 37C in PBS/0.2% Triton-X-100/10% DMSO/6% donkey serum, and incubated for 7 days in primary antibody (rabbit anti-cFos, 1:2000; Synaptic Systems #226-003) in PBS/0.2% Tween-20/10% DMSO/6% donkey serum and 10 ug/mL heparin. Following a wash step in PBS/0.2% Tween-20/heparin, brains were incubated in secondary antibody (donkey anti-rabbit Alexa Fluor 647, Thermo Fisher Scientific #A-31573) at 1:500 in PBS/0.2% Tween-20/10% DMSO/6% donkey serum/heparin for 7 days. Brains were washed again with PBS/0.2% Tween-20/heparin, then dehydrated a second time in a decreasing series of 20-100% methanol in PBS as above. The next day, brains were incubated in a solution of 66% dichloromethane/33% methanol for three hours, then two consecutive times in 100% DCM for 15 min each. After incubation in DCM, brains were transferred to a clean tube containing dibenzyl ether until imaging.

### Light sheet imaging

Cleared brain hemispheres were imaged in the sagittal orientation using a light sheet microscope (Ultramicroscope II, LaVision Biotec, Bielefeld, Germany) with an Olympus MVPLAPO 2X/0.5 objective and an Andor Zyla 5.5 camera. The dorsal surface of the brain was oriented closest to the light sheet, and images were collected using ImspectorPro software. Scans were performed using three angled light sheets, a light sheet numerical aperture of 0.026, sheet width of 100%, and Z step of 6 um. Brains were imaged in the 488 nm channel at 0.8x magnification for detection of autofluorescence as a single Z stack. The 647 nm channel was imaged at 2x magnification to image c-fos stained nuclei in 35-48 tiles (adjusted based on sample size). The field of view was cropped to 1000 × 1000 pixels, and stacks were acquired with a 10% overlap in X and Y. In a separate image processing step, tiled images were aligned using the Imaris Stitcher.

### ClearMap Analysis

Open source ClearMap 1.0 software was used to analyze c-Fos-positive cell counts across brain regions using previously described methods (16). The scripts were implemented in Python 2.7 on a Dell Precision Tower 5810 running Ubuntu 16.04 LTS. Cell detection parameters included cell shape parameter threshold set to 8 pixels in diameter, and Difference of Gaussian smoothing was performed using a kernel of 8 × 8 × 4 voxels. Detected cells were registered to the Allen Brain Atlas 25 µm map using 3D transformation between the 647 and 488 channels, and the 488 channel and the Allen Brain Atlas map, in order to compute c-fos-positive cell counts for each region.

### Functional network construction

We used the c-fos-positive cell counts to compute the between-subjects Pearson correlation coefficient *r* for all pairs of brain regions (110 regions) and defined it as our measure of functional connectivity. Because thresholding is arbitrary and may introduce biases such as false negatives and network density differences that skew group comparisons (17–19), and bias from data acquisition artifacts can cause false strong correlations, we chose not to apply thresholds. We constructed our networks as full correlation matrices, resulting in signed (positive and negative), undirected, weighted functional networks representing the neural state of each condition. Each node in the network was labeled with its anatomical group according to the Allen Brain Atlas.

### Network comparisons

To compare network properties among the four treatment groups (LD-abstinence, LD-reaccess, HD-abstinence, HD-reaccess), for each metric of interest we computed the empirical difference *D*^*emp*^ between groups and built a null model via a permutation procedure. First, we shuffled the condition labels among subjects without replacement, reconstructed the networks, and calculated the metrics and their condition difference *D*^*null*^ under the random null hypothesis. If the metric was community-based, we used the empirical community partitions in the null model networks. We built a null model distribution based on 10000 permutations. Finally, we calculated a p value for each comparison, defined as the proportion of the *D*^*null*^ distribution that can explain *D*^*emp*^, *p = (D*^*emp*^ *– D*^*null*^*) / N -1*, where *N* = 10000. The level of significance for each comparison was α = 0.05, and FDR correction of the p value was performed in nodal level comparisons.

### Community detection and global metrics

Many network properties arise from the configuration of nodes into complex network-wide interaction patterns, described as communities, clusters, or modules. To capture community structures, we used a pipeline largely developed by a previous study on multiscale hierarchical consensus clustering (HCC) (Jeub et al., 2018). We computed three global level metrics for each network: mean functional connectivity (mean *r*); anatomy-based modularity, which is the modularity quality function calculated using the anatomical groups for the community partition; and community-based modularity, calculated from the finest partition of the HCC procedure. We compared these metrics among the four treatment groups. To identify changes in the position of importance of individual brain regions among treatment conditions, and to examine how interactivity with other regions contributed to these changes, we computed two metrics to capture the position of a node within its network: within-community strength (wcs) and the diversity coefficient (h). For details of the community detection procedure and network metric calculations, see *Supplemental Methods*.

All network analysis was performed using R 4.0.2 (R Core Team, 2021) in Rstudio1.3.959 (RStudio Team, 2020), and the packages igraph (Csardi & Nepusz, 2006), boot (Canty & Ripley, 2020), data.tree (Glur, 2020), statGraph (da Costa et al., 2020), ggplot2 (Wickham, 2016), ggpubr (Kassambara, 2020), pracma (Borchers, 2021), psych (Revelle, 2021) and custom codes translated from Matlab from the Brain Connectivity Toolbox https://sites.google.com/site/bctnet/. All the codes used in this study can be found at https://github.com/coelhocao/Brain_Network_analysis.

### Chemogenetic silencing of cortical amygdala neurons

8-week-old mice were stereotaxically injected with AAV8-hSyn-hM4Di-mCherry or control AAV8-hSyn-mCherry virus in bilateral COA (coordinates: -1.7 AP, +-2.8 ML, -5.9 DV). Because c-fos counts were elevated in CIE mice in both anterior and posterior COA, we chose relatively central injection coordinates that would allow viral diffusion to both subregions (20). Following a 2-week surgical recovery period, mice underwent 4-5 weeks of baseline drinking and 4 cycles of CIE as described above. Beginning in test 3, mice received daily i.p. saline injections 30 minutes prior to voluntary drinking sessions. On drinking day 3 of test 4, corresponding to the day of brain tissue harvest for iDISCO in the previous cohort, mice received an i.p. injection of 3 mg/kg CNO 30 minutes prior to drinking. The following week, sucrose preference tests were performed. Mice were given 2h access to either water 5% (w/v) sucrose solution for habituation on the first day. On the following two days, mice received an i.p. injection of either saline or 3 mg/kg CNO, and 30 min later were given 1h access to sucrose or water. One week after sucrose testing, locomotor activity was tested 30 min following a 3 mg/kg CNO injection using a SuperFlex boxes (Omnitech Electronics, Accuscan, Columbus, OH). At the completion of all behavioral experiments, mice were perfused and brains extracted for verification of viral placement.

### Data analysis

C-fos count data were analyzed by 2-way ANOVA and Tukey’s post-hoc tests using R software. The level of significance for each comparison was α = 0.05, and an FDR of 5% was applied to correct for multiple comparisons. For drinking experiments, fluid consumption was determined by weighing bottles before and after drinking sessions. An empty cage mounted with a bottle containing the indicated fluid (alcohol, water, or sucrose) was used to estimate weight loss due to fluid drip, and drinking values were corrected accordingly. Consumption was expressed as g/kg body weight (adjusted for respective fluid density). Drinking data and behavioral data were analyzed in Prism using 2-way or 3-way ANOVA as appropriate followed by Tukey’s post hoc testing. Alpha was set at 0.05 for all comparisons.

## Results

### Chronic intermittent ethanol vapor increased voluntary ethanol drinking

To explore the effects of alcohol dependence on brain-wide patterns of c-fos activation, we used the CIE/FSS paradigm, a well-characterized model of alcohol dependence and stress-induced escalation of drinking. After establishing stable baseline drinking, mice underwent four cycles of CIE vapor or air exposure with alternating weeks of alcohol drinking, and a subset was subjected to 10 minutes of forced swim stress 4 hours prior to each voluntary drinking session (Figure 1A). Consistent with the established model, CIE mice displayed an increase in voluntary alcohol drinking that was statistically significant in tests 3 and 4 (Fig 1B, main effect of group [F(3,44)=19.53, p<0.0001]; group x week interaction [F(12,176)=12.12, p<0.0001]; test 3, CIE vs. AIR, p=0.0001). This effect was enhanced when CIE mice underwent FSS prior to drinking, whereas FSS had no effect on drinking in controls (test 4, CIE vs. CIE FSS, p<0.05; AIR vs AIR FSS, p=0.78). BECs did not differ between CIE and CIE+FSS mice during any week of vapor exposure (Fig 1C, CIE vs. CIE FSS [F (1, 25) = 0.002456, p=0.96]). On the day of brain tissue harvest, one group of mice was sacrificed without alcohol drinking (abstinence), while the remaining mice were given 1h of alcohol reaccess prior to sacrifice. As expected, alcohol consumption in the subset of mice that drank on the day of sacrifice was representative of the cohort as a whole, where CIE mice drank more than both AIR and AIR+FSS groups, and CIE+FSS tended to further escalate drinking (Fig 1D, significant effect of group [F (3, 44) = 34.82, p<0.0001]; CIE vs. AIR, p<0.0001; CIE vs. AIR+FSS, p=0.0001; CIE vs. CIE+FSS, p=0.004).

**Figure 1.**
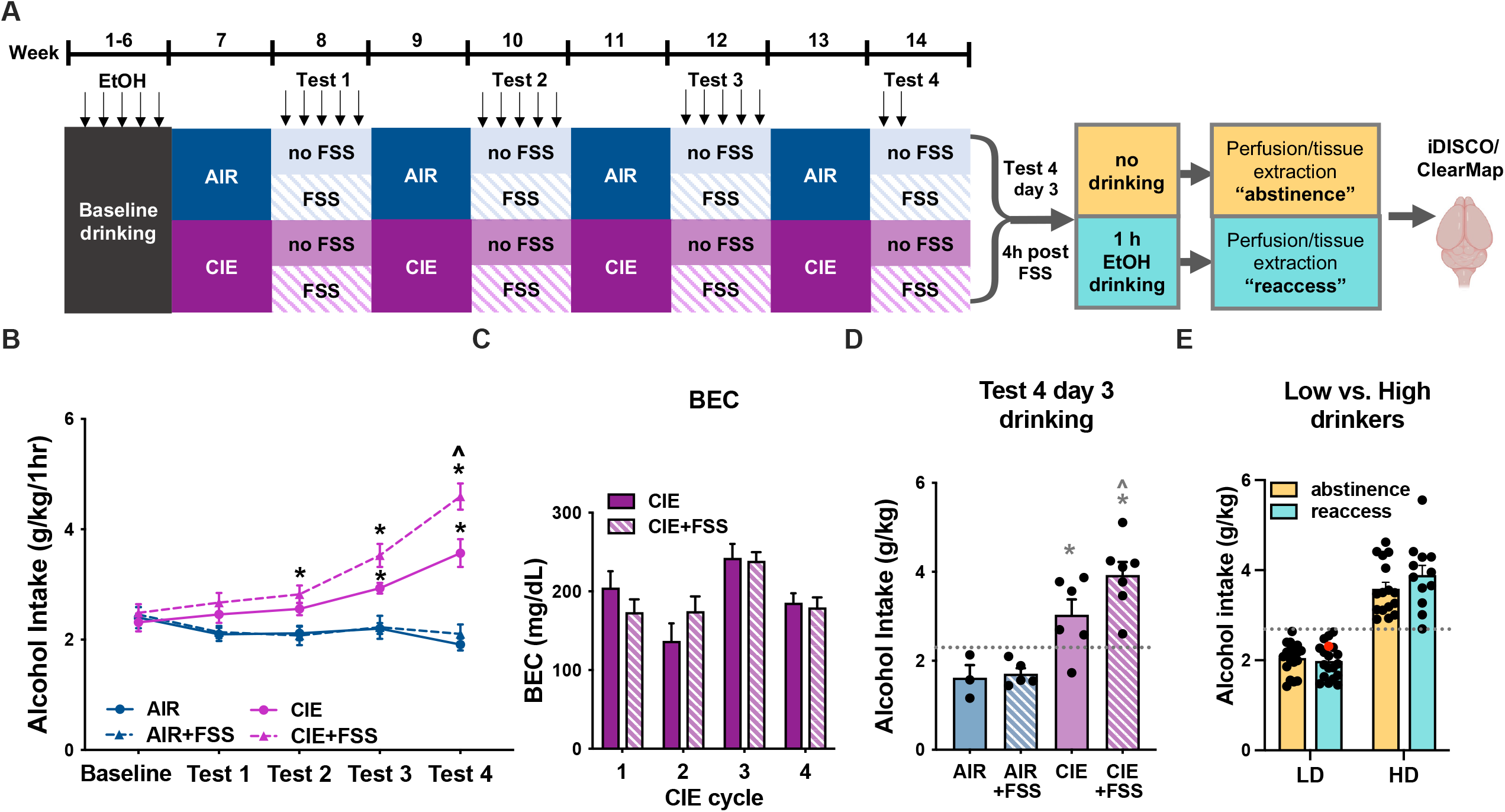
A, Timeline of chronic intermittent ethanol (CIE) and forced swim stress (FSS) treatment. Following 6 weeks of baseline drinking with 15% EtOH for 1h daily, mice underwent four cycles of AIR/CIE. On alternating weeks, mice had 1h daily access to alcohol. A subset of CIE and AIR mice underwent FSS 4h prior to each drinking session. On day 3 of test 4, a subset of mice in each group was sacrificed during abstinence, with the remaining sacrificed following a 1h drinking session (“reaccess”). B, CIE mice increased their voluntary alcohol intake across the four alcohol vapor exposure cycles. FSS further escalated drinking in CIE but not AIR mice (*, p<0.05 vs. AIR; ^, p<0.05 vs. CIE). C, Blood ethanol concentrations (BEC) during the four alcohol vapor exposure weeks. BECs did not differ between CIE and CIE+FSS mice. D, The subset of mice that drank on the final test day showed expected drinking behavior, where CIE increased drinking and FSS further escalated drinking in CIE mice (*,p<0.05 vs. AIR; ^, p<0.05 vs. CIE). E, For network analysis, mice were divided into groups of low and high drinkers (LD and HD, respectively) based on a cutoff of 2.7 g/kg averaged across tests 3 and 4, indicated by the dotted gray line. Only one CIE mouse, highlighted in red, showed consumption below this cutoff. During the final reaccess drinking session, alcohol intake was significantly higher in HD vs. LD mice (p<0.05), with the group separation indicated by the dotted gray line in 1D.

### Alcohol withdrawal causes widespread neuronal activation in CIE mice

We next used the iDISCO protocol to perform whole-brain c-fos immunolabeling and clearing of hemisected brains, followed by light sheet imaging and ClearMap automated analysis. ClearMap is a well-validated method for quantifying the number of c-fos-positive cells in brain regions registered to the Allen Brain Atlas. Figure S1A-D shows representative images of c-fos-stained nuclei in cleared brain tissue, and Figure S1E-F shows associated c-fos counts from selected brain regions. A complete list of included brain regions and raw c-fos counts from individual subjects is included in Tables S3 and S4. We first assessed the impact of CIE and FSS on neuronal activation at the abstinence time point by conducting two-way ANOVAs on raw c-fos counts for individual brain regions. Using a false discovery rate of 5% to account for multiple comparisons, regions with raw p values <0.0063 were considered statistically significant. CIE produced widespread increases in c-fos expression, with the effect size reaching significance in 40 regions (Figure S2, main effect of CIE): lzHY, BA, PAA, SI, COA, PA, SOC, PVZ, NLOT, BST, VIS, MEA, ARH, MRN, MA, BMA, PRN, PIR, STR, PT, DORsm, AAA, CA2, PB, SPF, IA, CA3, SCm, PV, STRd, CUN, CA1, PVH, PAG, STRv, OT, ENT, SCs, PPN, and sAMY. FSS increased c-fos in four regions, comprising the primary sensory cortices (SS, AUD, and VIS) and the sensory-related superior colliculus (SCs). While FSS showed a tendency to augment the increase in c-fos driven by CIE, no FSS x CIE interaction was detected for any region. A complete list of statistical results including post-hoc test results for each region is available in Table S5.

### Alcohol reaccess drinking differentially affects c-fos in high and low drinkers

A low n (n=3) in the AIR reaccess group due to a tissue processing error precluded a full group analysis for the reaccess condition. Because we detected minimal effects of FSS on c-fos counts, and network analyses are highly susceptible to errors driven by a small sample size, we collapsed the four groups (AIR, AIR FSS, CIE, and CIE FSS) into two groups comprising high and low alcohol drinkers (HD and LD) for subsequent analyses. Drinking values from tests 3 and 4 were averaged to minimize the influence of daily variability in drinking, and mice averaging at least 2.7 g/kg/h were designated as HD mice; this group comprised CIE and CIE FSS mice, with only one CIE mouse reassigned to the LD group (Figure 1E). We then explored the patterns of c-fos activation associated with differential drinking behavior. Two-way ANOVAs were conducted on raw c-fos counts for individual brain regions, with drinking history and reaccess as the independent variables. Using a false discovery rate of 5% to account for multiple comparisons, regions with raw p values <0.0082 were considered statistically significant. Reaccess alcohol drinking had a main effect to increase c-fos in STRd, while it decreased c-fos in four regions: SCm, SCs, VIS, and GENv (Figure 2). There were 13 regions with a significant effect of drinking history (HD vs. LD): AAA, BA, IA, STR, STRd, TEa, VIS, CLA, COA, NLOT, PAA, lzHY, PIR. In each of these regions, HD mice had higher c-fos than LD mice, driven primarily by the elevated c-fos during abstinence. The most prominent result of this analysis was observed in the interaction between reaccess drinking and drinking history: 34 regions showed differential effects of reaccess drinking in LD vs. HD mice, where drinking increased c-fos in LD mice but suppressed c-fos in HD mice. These regions included CA1, CA2, CA3, DG, BST, BA, MEA, MS, SI, VIS, BMA, PA, SPF, DORsm, CUN, MRN, PAG, PPN, RN, SCm, SNc, SNr, COA, NLOT, PAA, ARH, DMH, lzHY, mzHY, PV, PVZ, VMH, PB, and PRN (Figure 2). Statistical results including post-hoc test results for each region are included in Table S5.

**Figure 2.**
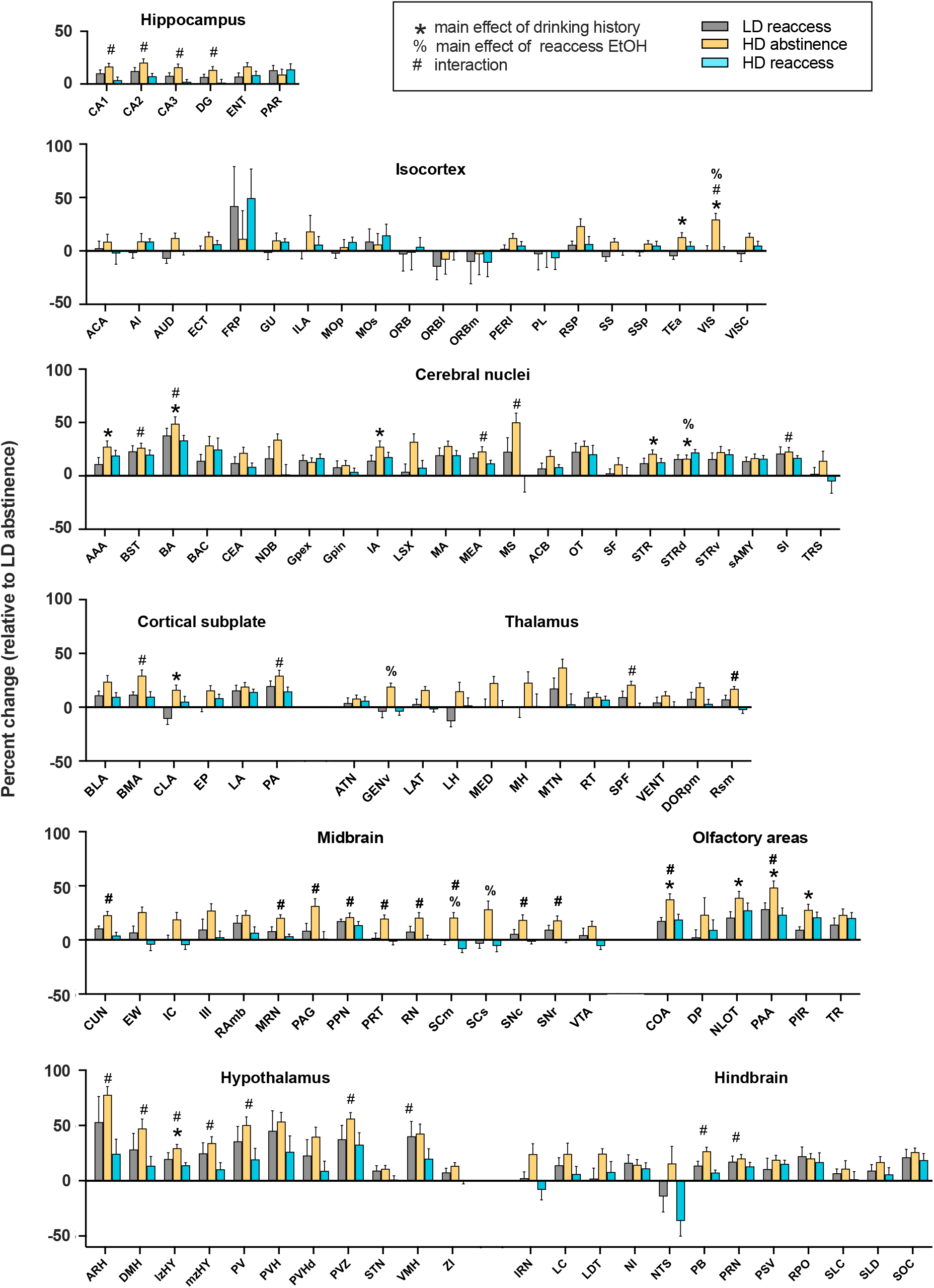
A, Relative c-fos expression in high- (HD) and low-drinking (LD) mice during acute (24h) alcohol abstinence and following a 1h alcohol reaccess period. C-fos-positive cell counts for each region are represented as the percent change relative to LD mice during abstinence. Regions are grouped by anatomical subdivision according to the Allen Brain Atlas. * denotes a significant main effect of drinking history; % denotes a significant main effect of reaccess EtOH drinking; and # denotes a significant interaction between drinking history and alcohol reaccess (2-way ANOVA with FDR correction for multiple comparisons). Post-hoc test p-values are available in Table S5.

### Chronic alcohol and reaccess drinking reorganize functional brain networks

We next performed a network analysis to assess how a history of high-level alcohol exposure and/or reaccess drinking changed the patterns of coactivation among brain regions. We built fully connected, undirected, signed, weighted networks for the LD abstinence, LD reaccess, HD abstinence, and HD reaccess conditions, generated from between-subject interregional correlations (Figure 3A-D). The hierarchical consensus clustering procedure revealed a nested community structure in which the networks were divisible into smaller clusters of increasingly strong interregional correlations for up to 6 levels (Figure 3E-L). This hierarchical organization was statistically above chance and consistent with observations in anatomical human and rodent networks (Jeub et al., 2018). The network communities for each condition are shown in Figure 4.

**Figure 3.**
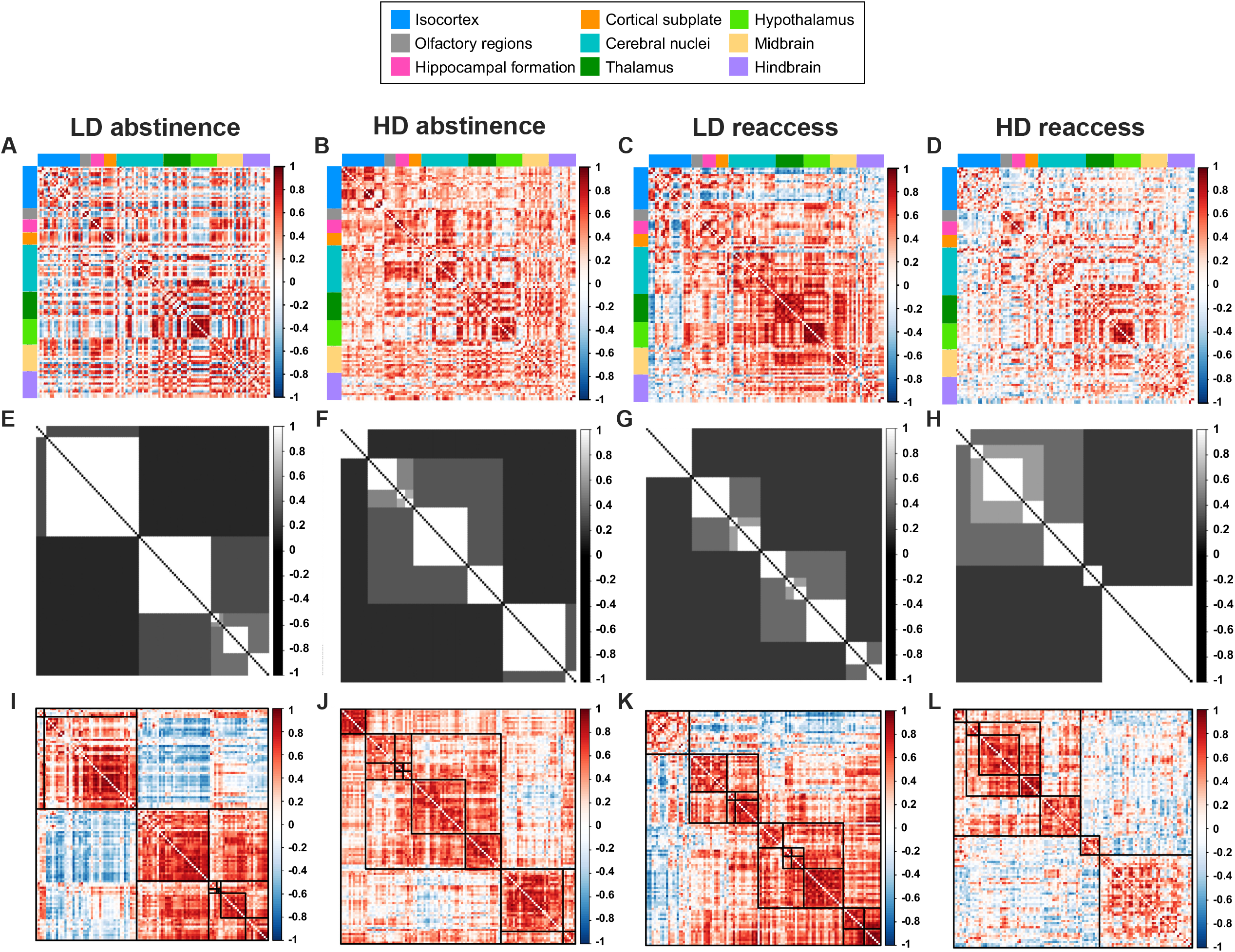
A-D, Pearson correlation matrices showing interregional correlations for the 110 brain regions included in the network analysis for low-drinking (LD) and high-drinking (HD) mice during alcohol abstinence and following alcohol reaccess. Regions are grouped by anatomical division according to the Allen Brain Atlas. E-H, Hierarchical consensus clustering (HCC) plots illustrating the community partitions of highly correlated regions. I-L, Correlation matrices derived from the HCC procedure illustrating the clustering of strongly correlated regions, denoted by darker shades of red.

**Figure 4.**
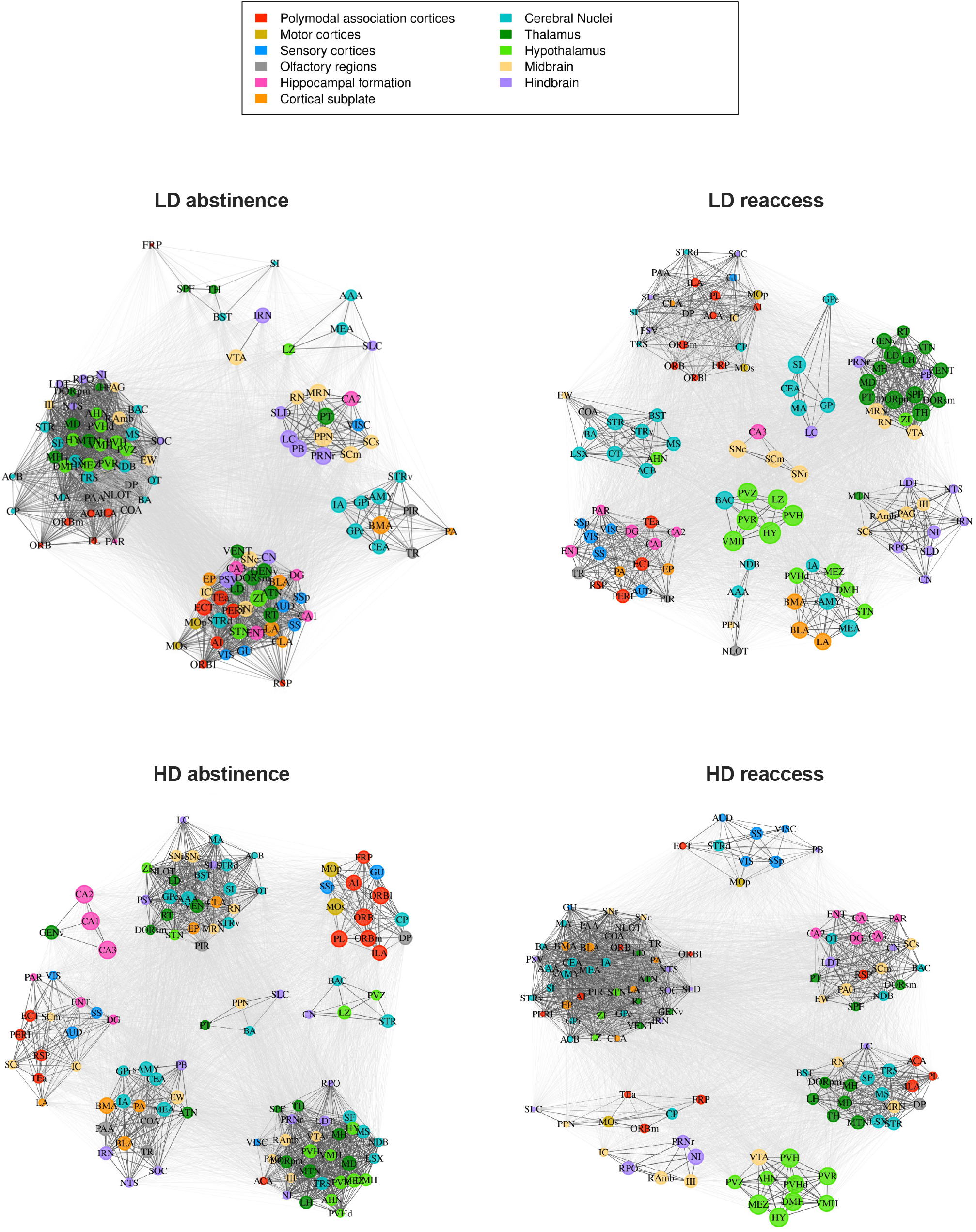
Network communities for high-drinking (HD) and low-drinking (LD) mice in the abstinence and reaccess conditions. Regions are color-coded according to anatomical groups defined by the Allen Brain Atlas. The size of each node in a community represents the within-community strength, whereas the strength of individual connections is represented by the edge length, i.e., the distance between nodes.

We next computed the adjusted mutual information (AMI) between the HCC community partitions and the anatomical group partitions as defined by the Allen Brain Atlas (Figure S3). AMI indicated a high degree of overlap between HCC and anatomical clusterings for all four groups. This finding is represented in Figure 4, where communities are largely composed of multiple regions from the same anatomical group, specifically the polymodal association cortices, hippocampal formation, and cerebral, thalamic, and hypothalamic nuclei. This partial correspondence between the functional communities and anatomical groups was observed in all conditions and indicates that the HCC communities may be capturing interaction patterns of coordinated regions belonging to both the same and different anatomical systems, revealing information exchange across systems.

For each network, we calculated the mean functional connectivity (FC), anatomical modularity, and hierarchical consensus clustering (HCC) modularity (Table S1). Whereas anatomical modularity is calculated given the anatomical groups from the Allen Brain Atlas, HCC modularity is computed using the communities derived from HCC clustering. Because only a single network was generated for each condition, we made statistical comparisons between group pairs by computing the empirical difference between groups and building a null model using a permutation procedure (Table S2). Mean FC was not statistically different among the groups (Table S2). However, there was a significantly lower anatomical modularity in the HD-abstinence network compared to HD-reaccess network, indicating that reaccess alcohol drinking increased the interactivity of intra-anatomical groups in HD mice. This effect of reaccess alcohol on anatomical modularity was not observed in LD mice. We also observed a higher HCC modularity in the HD-reaccess network compared with both HD-abstinence and LD-reaccess networks (Table S2). These differences are visualized in Figure 3A, where the HD-reaccess network has a larger number of negative correlations (shown in blue) than the HD-abstinence and LD-reaccess groups, resulting in more segregated communities.

### Chronic alcohol and reaccess drinking alter regional connectivity metrics

One of the goals of our study was to identify brain regions that significantly change their interaction patterns following chronic alcohol exposure, and to understand how such changes reflect the community structure and overall change in brain state. After performing hierarchical consensus clustering and obtaining the functional communities, we computed the within-community strength (wcs) and diversity coefficient (h) for each region given its community allegiance and compared these metrics among conditions in a pairwise manner. Wcs captures the strength of a region’s connectivity within its assigned community, whereas h captures the strength of its connectivity to other communities. Figure 5 shows the regions that were statistically different in these two-by-two comparisons (Fig 5A-F); a complete table of significant results is available in Table S6. Overall, relatively few regions showed significant changes in wcs between conditions. The substantia innomminata (SI) was significant in two comparisons; wcs was higher in HD relative to LD mice in the abstinence condition (Figure 5A, p<0.05), and reaccess drinking increased wcs of the SI in LD mice (Figure 5B, p<0.05). Interestingly, the SI also showed different community associations in HD and LD mice (Figure 4); in HD mice, it was part of a large densely connected community spanning multiple anatomical groups, whereas in LD mice, regardless of reaccess alcohol drinking, it was part of a smaller cluster composed mainly of other cerebral nuclei. Thalamic regions also showed significant wcs differences across conditions, although this was not specific to any one subregion. Reaccess drinking increased the wcs in several thalamic subregions in LD mice (Figure 5B), whereas in HD mice, alcohol reaccess reduced wcs in many thalamic subregions (Figure 5C).

**Figure 5.**
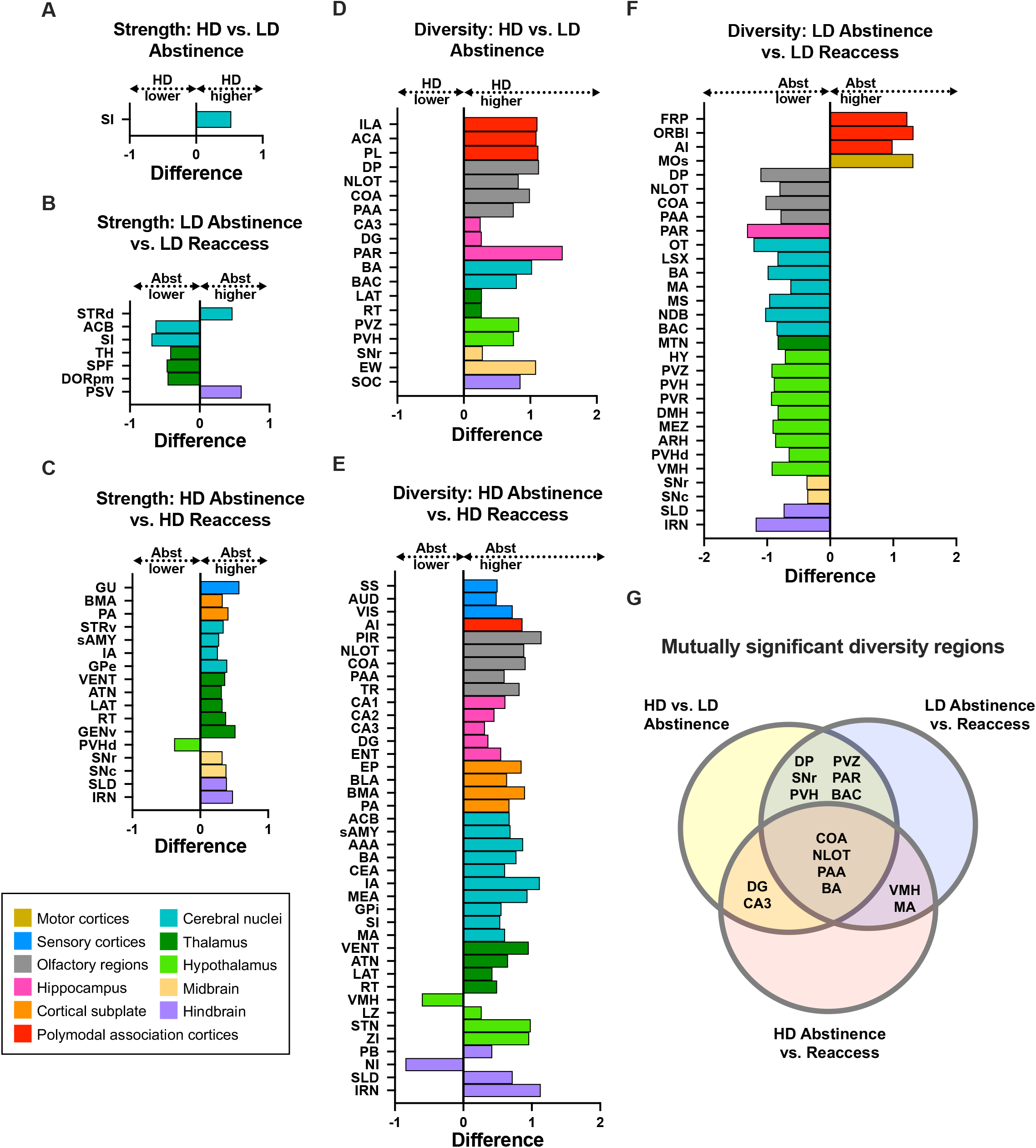
Significantly different brain regions for within-community strength and diversity coefficient comparisons. A-C, Differences in within-community strength between high drinkers (HD) and low drinkers (LD) during abstinence (A), LD abstinence and reaccess (B), and HD abstinence and reaccess (C). D-F, Differences in regional diversity coefficients between HD and LD abstinence (D), HD abstinence and reaccess (E), and LD abstinence and reaccess (F). G, Regions with significantly different diversity coefficients shared across multiple 2-condition comparisons.

We observed a similar pattern of results for the diversity coefficient (Figure 5D-F). Broadly, diversity coefficients for many regions were higher in HD than LD mice during abstinence (Figure 5D). Reaccess alcohol intake had opposite effects in HD and LD mice, reducing diversity across many regions in HD mice (Figure 5F), but increasing it in LD mice (Figure 5E). To identify regions of high importance, whose diversity coefficient changed across multiple conditions, we represented the mutually significant regions in a Venn diagram (Figure 5G). We were particularly interested in regions that showed an elevated diversity coefficient in HD mice during abstinence that was reduced by reaccess drinking; activity in these regions could be contributing to the high levels of c-fos activation occurring across communities during the abstinence condition, which may contribute to the elevated drive to drink alcohol in these subjects. Regions fitting these criteria included the DG and CA3 in the hippocampus, as well as four regions that were significant across all three comparisons: three olfactory regions—cortical amygdala (COA), nucleus of the lateral olfactory tract (NLOT), and piriform-amygdalar area (PAA)—as well as the bed nucleus of the accessory olfactory tract (BA), an anatomically adjacent structure with vomeronasal functions (21). Interestingly, the diversity coefficient of these four regions was reduced by reaccess alcohol in LD mice, suggesting that acute modulation of these areas by alcohol may result in long-term changes in functional connectivity with repeated high-level exposure.

### Inhibition of cortical amygdala neurons reduces drinking in CIE mice

Our network analysis identified several regions with higher diversity in the HD abstinence condition compared to LD-abstinence and HD-reaccess, suggesting these regions may have an important influence on cross-community network activity during abstinence and contribute to the heightened brain-wide activity that we suspect drives relapse drinking. We chose to test this hypothesis in one region, the COA, a chemosensory brain region known to mediate odor influences on motivated behaviors. The COA exhibits reciprocal connections with a number of subcortical areas implicated in alcohol dependence—basal forebrain, BNST, hypothalamus, thalamus, and brainstem—and also projects densely to CeA (22). Using designer receptors exclusively activated by designer drugs (DREADDs) (23), we inhibited this brain region during an alcohol reaccess session. Mice were stereotaxically injected with an AAV expressing either hm4di-mcherry or mcherry control virus in bilateral COA, and following a recovery period, underwent 4-5 weeks of baseline drinking followed by four cycles of CIE (Figure 6A). Mice received saline injections 30 minutes prior to each drinking session beginning in test 3, which is reflected in a reduction in drinking in both AIR and CIE mice at this time point (Figure 6G). However, this did not obscure an effect of CIE, which significantly increased drinking in tests 3 and 4 (Figure 6G). Following the fourth week of vapor exposure during drinking test 4, mice were given 3 mg/kg CNO 30 minutes prior to drinking during the third daily drinking session. Because individual mice exhibit high variability in day-to-day drinking, alcohol intake following CNO was compared to the average consumption on the three surrounding days when mice received saline injections (Figure 6I). CNO reduced drinking selectively in CIE mice expressing the hm4di virus (3-way-ANOVA, trend for CNO x CIE x virus interaction, p=0.056, CNO x CIE interaction, p=0.0024, CIE hm4di CNO vs. CIE hm4di saline, p=0.0002, CIE mcherry CNO vs. CIE mcherry saline, p=0.24). We also computed the percent change in drinking for individual mice and detected a significant effect of CNO in CIE hm4di mice only (one-sample t-test, CIE hm4di, p=0.0005, all other groups, p>0.18, Figure 6J). To determine if the reduction in drinking with CNO in CIE hm4di mice was selective to alcohol, we performed a test of sucrose consumption the following week. There was no effect of CNO on sucrose consumption, water consumption, or sucrose preference (Figure 6K-M). However, we did detect a main effect of hm4di virus to reduce sucrose consumption (3-way-ANOVA, p=0.03, Figure 6K). We also tested whether the reduction in alcohol consumption could be due to an overall reduction in locomotor activity; no differences in distance traveled or mean velocity were detected in a 30-minute test of locomotion performed 30 minutes following a CNO injection (Figure 6N,O).

**Figure 6.**
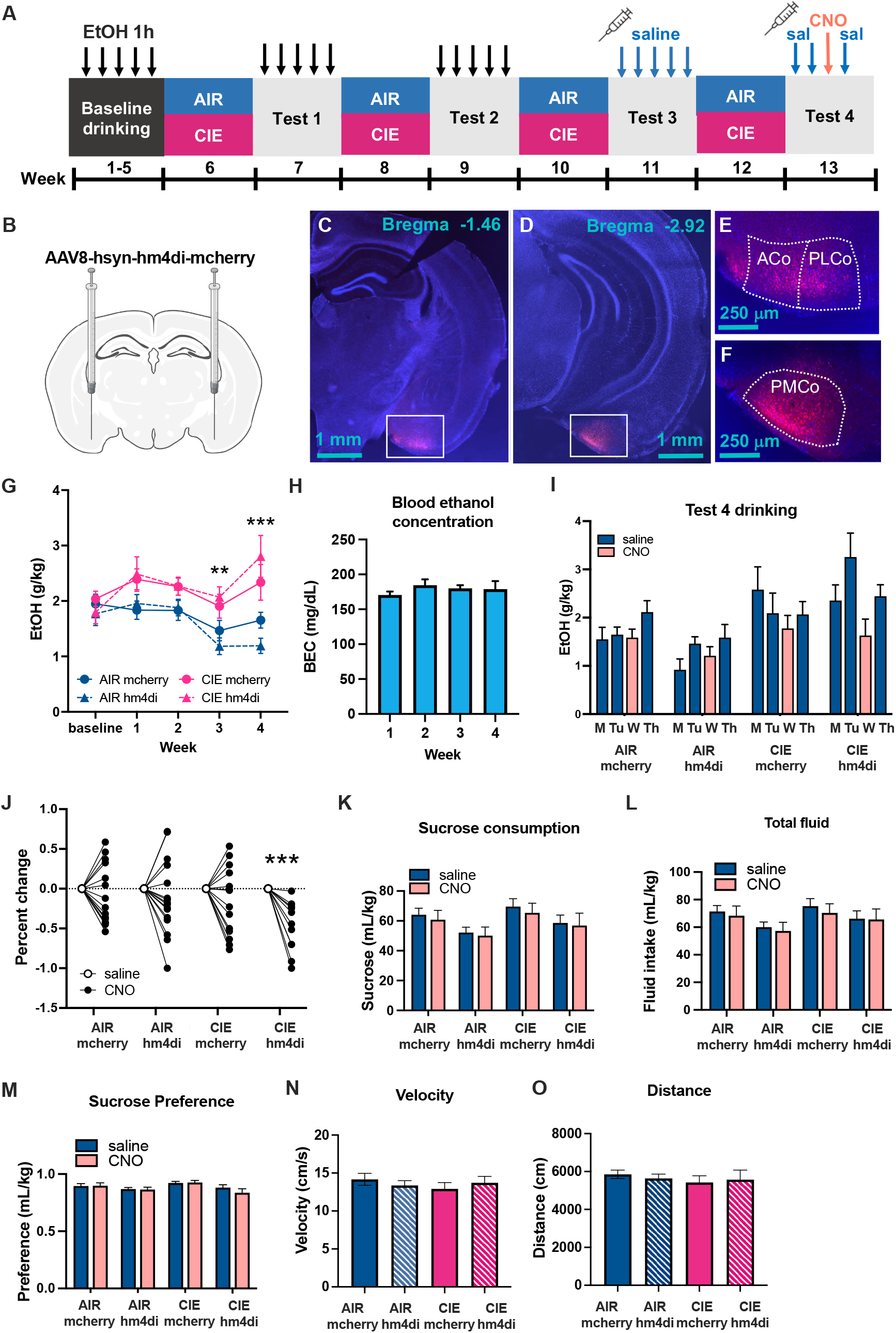
A, Experimental Timeline. Following 5 weeks of baseline drinking of 15% EtOH daily for 1h, mice underwent 4 cycles of AIR/CIE vapor with alternating weeks of daily 1h alcohol access. During drinking tests 3 and 4, mice received daily saline injections 30 min prior to drinking. On day 3 of test 4, mice were injected with 3 mg/kg CNO instead of saline. B, Viral injection schematic. C-F, Representative images of viral placements in cortical amygdala showing a subject with anterior placement (C,E) and a subject with posterior placement (D,F). G, Alcohol intake was higher in CIE than AIR mice during test weeks 3 and 4, with no effect of mcherry vs. hm4di genotype. H, BECs were relatively stable in CIE mice across the four vapor exposure weeks. I, Alcohol drinking during test 4. Mice received saline on days 1, 2, and 4, and CNO on day 3. J, Data from individual mice showing the percent change in drinking following CNO administration relative to the average of saline days. CNO reduced drinking selectively in CIE mice expressing hm4di virus (one-sample t-test, p<0.001). K-M, Sucrose consumption (H), total fluid consumption (I), and sucrose preference (J) were unaffected by CNO. We detected a main effect of hm4di virus to reduce sucrose drinking independent of CIE or CNO treatment (p<0.05). N-O, Locomotor activity measured via distance traveled (K) and mean velocity (L) did not differ based on virus or CIE exposure after CNO administration.

## Discussion

Using whole-brain c-fos mapping and network analysis, we demonstrated that acute (24h) alcohol abstinence in high-drinking, CIE- and CIE/FSS-exposed mice produces widespread neuronal activation and changes in network community architecture marked by increased cross-module communication of multiple brain regions. Alcohol reaccess drinking suppressed this increase in neuronal activation and reversed the abstinence-induced changes in coactivation, increasing brain modularity. We identified several olfactory brain regions, including the COA, as potentially important drivers of cross-community neural activation during withdrawal. Gi-DREADD-mediated inhibition of the COA reduced voluntary drinking during acute abstinence in CIE mice, validating our network approach and suggesting that the COA, and other olfactory-associated brain regions, may be novel regions of importance for alcohol dependence.

Prior studies have assessed c-fos activation following acute and chronic alcohol intake, but this is the first to assess global c-fos patterns during acute alcohol abstinence and following alcohol reaccess in the CIE model. We identified a large number of activated brain regions during alcohol abstinence not restricted to those previously implicated in alcohol drinking. Chronic alcohol-induced molecular and cellular neuroadaptations, such as reduced inhibitory neurotransmission via GABAa receptors and increased excitatory glutamatergic transmission (24), produce a state of hyperexcitability during withdrawal. This is reflected in elevated c-fos, and given that alcohol’s pharmacological effects extend to the whole brain, the global nature of these changes is unsurprising. Upon re-exposure to alcohol, binge-level alcohol consumption in high-drinking mice suppressed the withdrawal-induced increase in c-fos, consistent with alcohol’s depressant effects at high concentrations. A recent study examining the time course of c-fos expression post CIE exposure made similar observations, albeit in a more restricted set of brain regions (25). This study showed reduced c-fos expression during acute alcohol vapor-induced intoxication, but increased c-fos across most regions at 26h of withdrawal, supporting the notion that acute abstinence represents a state of global hyperexcitability in dependent mice and that c-fos reliably indicates these changes.

The interaction between drinking history (HD vs LD) and reaccess alcohol drinking was significant for a large number of brain regions. Whereas reaccess drinking suppressed c-fos in HD mice, it increased c-fos in many of the same brain areas in LD mice. Consistent with the latter, studies of alcohol-induced IEG expression in rats and mice, focusing on addiction-related brain regions, show nearly universal increases in c-fos following acute i.p. injection or voluntary consumption of 1-4 g/kg alcohol in naive animals or following short-term repeated exposure. The differential response of HD and LD mice to acute alcohol may be due in part to differing alcohol doses, as HD mice consumed approximately double the volume of ethanol as LD mice. However, this difference might also reflect the neuroplasticity resulting from chronic high-level alcohol exposure that changes its pharmacological outcome. Interestingly, although LD and HD mice had different baseline c-fos values during abstinence, and consumed very different amounts of alcohol, c-fos counts post-drinking were similar for most regions. This suggests that mice drink until reaching a similar homeostatic point, and allostatic mechanisms in dependent mice increase the amount of alcohol required to attain homeostasis. The regions affected include many already implicated in drinking, such as the extended amygdala and lateral hypothalamus, as well as many understudied regions such as the substantia innominata.

One goal of our study was to gain insight into how FSS promotes increased drinking in CIE mice. We found that 4 hours post FSS, there was increased activation of sensory-related systems in both AIR and CIE mice, specifically in the auditory, somatosensory, and visual cortices, and in the sensory-related superior colliculus. Sensory hyperreactivity is a known consequence of acute ethanol withdrawal (26), and FSS may exacerbate this state in CIE mice by further disrupting sensory processing. Although we did not detect a statistical interaction between FSS and CIE for any region, CIE mice that underwent FSS showed a tendency toward further elevation of c-fos across many brain regions during abstinence, in particular the cerebral nuclei, hippocampus, and olfactory regions. Consistent with this, a recent whole-brain c-fos mapping study in mice found that acute stress increased c-fos across a majority of brain regions examined, supporting the idea that stress promotes widespread increases in activation (27). This trend for increased c-fos following FSS was not observed in AIR mice, coinciding with their lack of escalation in drinking. Mice exhibit hormonal and behavioral habituation to repeat stressors, including FSS (28), suggesting AIR mice may have adapted to FSS over time. CIE mice show an elevated corticosterone response to a single FSS (29), and may show differential adaptation to repeat FSS due to alcohol dysregulation of stress systems (30), although this was not explicitly assessed. Given the timing of tissue collection—four hours post stressor—differences in stress recovery between AIR and CIE may be particularly important, and this is an interesting area for future study.

When we analyzed how these global c-fos changes translated into network function changes, several patterns emerged. During abstinence, there were no differences between LD and HD mice in general network features such as mean functional connectivity or modularity; however, the community configurations differed, and many regions in the HD abstinence group had an increased diversity coefficient, indicating substantial coactivation with regions outside of their assigned communities. Following reaccess drinking, network properties were significantly altered in HD but not LD mice. Alcohol reaccess increased HCC modularity in HD mice, indicating that it increased the segregation of the communities identified by HCC, and also increased anatomical modularity, indicating greater interactivity of intra-anatomical groups. Similarly, alcohol reaccess reduced the diversity coefficients in many regions in HD mice, indicating lower inter-community connectivity, consistent with the increase in modularity.

In previous work in CIE mice, assessed after one week of alcohol abstinence, reduced modularity was found to be the major hallmark of abstinence (15), and this was also observed for several other drugs of abuse (31). Modular organization, where neural elements form strong within-module connections and weak inter-module connections, is a feature of complex networks that supports specialized processing (32). We did not detect a reduction in overall brain modularity during abstinence, but the increase in inter-module connectivity of many regions suggests similar functional changes at the circuit level in our model. Increased modularity following alcohol was at least partly due to an increase in negative correlations resulting from alcohol inhibition of circuit activity. Acute withdrawal may produce nonspecific activation of extraneous circuits, in essence increasing the noise in the system, and drinking may increase modularity by suppressing this inappropriate circuit activity. The alcohol-dependent brain may therefore function more efficiently after alcohol reaccess, contributing to the drive to consume alcohol.

Our network findings echo those of two recent human fMRI studies, where alcohol-dependent subjects showed changes in the network community structure at baseline, and relapse caused the community configuration to become more similar to that of control subjects. In one study, abstinent drinkers displayed increased global cross-community interaction of the vmPFC, particularly with the visual-cortex containing community, and this difference was reversed following relapse (33). Similarly, we observed a significantly elevated diversity coefficient in abstinent HD mice in PL, ILA, and ACA, rodent correlates of human PFC (34); while reaccess tended to attenuate these diversity measures in high drinkers, this finding did not reach significance. In another human imaging study, basal ganglia and thalamic nodes had different community associations in heavy drinkers compared to controls during abstinence, and returned to similar configurations as controls following relapse (35). However, modularity was not different in drinkers relative to controls either before or after relapse. In line with our findings, these two studies suggest that in alcohol-dependent human subjects, drinking normalizes aspects of network function that have likely been disrupted by chronic alcohol use, although baseline genetic contributions to the observed phenotypes cannot be ruled out.

Network analysis pointed to a cluster of anatomically adjacent olfactory-related brain regions that may be particularly susceptible to modulation by both chronic alcohol and reaccess drinking. These included the COA, PAA, NLOT, and BA, which showed increased c-fos activation after reaccess alcohol in LD mice, and increased c-fos during abstinence in HD mice that was reversed by reaccess drinking. Similar patterns were observed for network diversity coefficients. One possible explanation for these effects is that chronic alcohol vapor inhalation damages peripheral olfactory pathways. While this has not been studied in the CIE model, evidence in rodents shows that acute alcohol inhalation causes olfactory neuronal death, as well as retrograde changes in the hippocampus, another high-diversity region in our study. Alcohol-induced olfactory changes are not specific to the vapor route of administration, however. Chronic alcohol drinking causes olfactory damage in both humans and rodents (36–38). In humans, alcohol has acute dose-dependent effects on olfaction, where low-dose drinking increases olfactory perception, and high-dose drinking blunts olfaction (39). Thus, alcohol can both acutely alter olfaction and lead to chronic changes in olfactory brain regions over time, even when consumed orally. Our DREADD experiment supports the role of cortical amygdala in withdrawal-induced drinking, and the other olfactory regions identified are predicted to have similar effects. These regions and their differential modulation by alcohol may serve as useful biomarkers for alcohol misuse. Interestingly, chronic stress also causes significant olfactory dysfunction in rodents, mediated in part by changes in the NLOT (40), suggesting that the olfactory cluster we identified may be an important locus for stress-alcohol interactions.

A key limitation of this study is the use of the collapsed dataset for the network analysis, which precluded assessing the effects of FSS on network properties in CIE mice. These groups showed significant differences in drinking behavior, and c-fos counts tended to be higher in CIE+FSS mice. It is unknown if this would translate into an exacerbation of the observed network changes in CIE+FSS mice compared to CIE (e.g., further reduced modularity), or if distinct CIE FSS networks would emerge. However, it is reassuring that a high-centrality target region identified using the collapsed dataset was validated in a CIE cohort that did not undergo FSS. A more general limitation of this study is the use of c-fos as a proxy for neuronal activity. Although widely implemented for this purpose, there are instances in which neuronal activation is independent of IEG induction—specifically, conditions of chronic activation or net synaptic inhibition (41), as well as instances of glial c-fos induction in neuroinflammatory states (42). That our c-fos measures are in agreement with other indicators of neuronal excitability during withdrawal (electrophysiological, etc.) suggests a neuronal component to our observations; given the neurotoxic and inflammatory effects of alcohol, a glial contribution is also possible. While our data demonstrate interesting brain-wide patterns, region-level exploration of specific cell types remains an important area for future study.

## Acknowledgements

This work was supported by the National Institutes of Health (NIH) National Institute of Alcohol Abuse and Alcoholism (NIAAA) grant 2U01AA020911-11 to T.L.K., and National Institute of Mental Health (NIMH) R01MH119421 and funding from the Liquor Control Board of Ontario (LCBO) to P.W.F. Some figures for this manuscript were created using BioRender.com. We thank Pablo Ariel, Ph.D. and the UNC Microscopy Sevices Laboratory for assistance with light-sheet imaging. The Microscopy Services Laboratory, Department of Pathology and Laboratory Medicine, is supported in part by P30 CA016086 Cancer Center Core Support Grant to the UNC Lineberger Comprehensive Cancer Center. Research reported in this publication was supported in part by the North Carolina Biotech Center Institutional Support Grant 2016-IDG-1016.

## Supplemental Figure Legends

Figure S1. A-D, Representative sagittal images of c-fos-stained brain tissue used for ClearMap automated cell detection in low-drinking (LD) and high-drinking (HD) mice during alcohol abstinence and following alcohol reaccess. Insets show higher magnification images of the dentate gyrus (DG, blue) and sensory-motor cortex related thalamus (DORsm, purple). E, Clearmap-generated cell counts for the DG for the four treatment groups. During abstinence, HD mice showed higher c-fos expression in the DG relative to LD mice (p<0.05). HD mice had lower c-fos expression in the DG following reaccess as compared to abstinence (p<0.05). F, Clearmap-generated cell counts for the DORsm for the four treatment groups. During abstinence, HD mice showed higher c-fos expression in the DORsm relative to LD mice (p<0.01). HD mice had lower c-fos expression in the DORsm following reaccess as compared to abstinence (p<0.001).

Figure S2. Relative c-fos expression in CIE and/or FSS-exposed mice during acute (24 h) ethanol abstinence. C-fos-positive cell counts are represented as the percent change relative to AIR mice for each region. Regions are grouped by anatomical subdivision according to the Allen Brain Atlas. * denotes a significant main effect of CIE; # denotes a significant main effect of FSS (2-way ANOVA with FDR correction for multiple comparisons). Post-hoc test p-values are available in Table S5.

Figure S3. A-D, The adjusted mutual information (AMI) between the partition distribution used in hierarchical consensus clustering (HCC) and the HCC finest partition (in black) or the anatomical groups partition (in blue) for low-drinking (LD) and high-drinking (HD) groups in the abstinence and reaccess conditions. The AMI relative to the finest partition peaked at around 0.8 for the γ values that provided partitions closer to the finest HCC partition. The γ values at which the HCC finest partition peaked coincided with that at which the AMI relative to the anatomical groups partition also peaked, though at an AMI roughly around 0.5. Since AMI is an index where a value of zero means independence between the two variables compared, these AMI values indicate that the HCC community partition had a partial agreement with the anatomical groups. E-H, The y(β) distribution for each group used to compute the partitions in A-D.

