## Supplemental Methods for "Alcohol dependence modifies brain networks activated during abstinence and reaccess: a c-fos-based analysis in mice"

**Community detection**

*Multi-resolution Modularity –* The modularity quality function measures the strength of ‘clustering’ in a network, given a community assignment vector $\vec{g}$, where g_i_ is the cluster containing the node *i*. Most community detection algorithms are based on modularity function optimization. The modularity quality function used in this study was built from the Reichart and Bornholdt (43) version of the modularity quality function,

$$Q\left( \vec{g},\gamma\right)=\sum_{i,j=1}^{n} \left( A_{ij}-{\gamma P}_{ij} \right)\delta\left( g_{i},g_{j} \right),$$

(1)

where A is the adjacency matrix, P is the expected adjacency matrix under a null model, γ is the resolution

pair parameter, and δ (g_i_, g_j_) is the community co-assignment matrix, where δ = 1 if g_i_ = g_j_, and δ = 0 otherwise. The resolution parameter influences the size of the clusters and can be tuned to encompass different scales. When applying the community detection algorithm in the empirical networks (see algorithm section below), we used the following typical null model (44),

$$P_{ij}=\frac{k_{i}k_{j}}{v},$$

(2)

where k is the sum of all correlations of a given node, and v is the sum of all k values. We use this null model when applying the community detection algorithm in the empirical networks.

In this study, we used a variant of the modularity quality function, Q*, previously observed to be well-suited for correlation-based networks (42),

$$Q^{*}=Q^{+} + \frac{v^{-}}{v^{-}v^{+}}Q^{-}$$

(3)

The modularity variant Q* assumes an unequal importance to the positive and negative correlations and provides the negative correlations a lower weight, proportional to their presence, such that if a network has equal numbers of positive and negative correlations, the contribution of positive correlations is twice that of negative.

*Community detection algorithm –* We used a Louvain-like algorithm for signed networks adapted from the community_louvain.m function in the Brain Connectivity Toolbox (<https://sites.google.com/site/bctnet/>). The Louvain algorithm is a heuristic based on maximizing the modularity quality function. It starts by selecting random nodes, grouping them locally into small communities if that increases modularity, then treating the small community as a node and repeating until there is no further increase in modularity (42,43).

*Resolution parameter –* The resolution parameter γ in (1) influences the null model P such that it tunes the size of the communities detected. It works as an inverted tuning knob in which increasing γ will tend to detect smaller communities. This effectively allows the detection of communities at different scales in the network, varying from a coarse scale with large communities (or no community at all) to the finest scale in which nodes are singleton communities. We adopted a multi-resolution approach that covers that range. However, when applying community detection until all nodes are singleton communities, certain correlations need values of γ scales of magnitude higher than most correlations to be separated, which can potentially make a linear or exponential selection of γ values to cover the whole range a costly approach. To circumvent this, we implemented a strategy used previously (44,45) to both determine the [γ_min_, γ_max_] range and sample meaningful γ values within this range (values that result in a change of community detection). Briefly, the [γ_min_, γ_max_] range was determined such that γ_min_ was the largest γ where no communities were detected, and γ_max_ was the smallest γ where all nodes are singleton communities. We calculated γ_max_ as the smallest γ such that A_ij_ – P_ij_ ≤ 0 for all i and j. As in Jeub et al., we determined γ_min_ iteratively by first estimating γ_min_ using a small sample of partitions (20) at γ = 1, then sampled a new set of partitions using γ = γ_min_ – ε, where ε is a small constant (1^-10^) that ensures γ_min_ is sub-optimal, then use the new sample to update γ_min_. This procedure was repeated until the new sample consisted of only trivial partitions (no communities; for more details, see Jeub et al., 2018).

To sample γ values within the [γ_min_, γ_max_] range, we split the contributions of node pairs to modularity into ferromagnetic correlations E^+^(γ) = {(i, j) | i ≠ j, A_ij_ − γP_ij_ > 0} and antiferromagnetic correlations E^–^(γ) = {(i, j) | i ≠ j, A_ij_ − γP_ij_ < 0}. Note that changing the γ value will change the correlations that become antiferromagnetic. As proposed by Jeub, we used the relative magnitude of the antiferromagnetic correlations as a measure of scales,

$$\beta\left( \gamma\right)=\frac{\sum_{{}_{\left( i,j \right)\in E^{-}\left( \gamma\right)}} \left| A_{ij}-\gamma P_{ij} \right|}{\sum_{{}_{\left( i,j \right),i\neq j}} \left▒ A_{ij}-\gamma P_{ij} \right|}$$

(4)

Note that when γ ≤ 0, β(γ) = 0, and when γ ≥ γ_max_, β(γ) = 1, and β increases monotonically for 0 ≤ γ ≤ γ_max_, that is, γ increases as a continuous variable, but β increases as if it were a discrete variable, changing when the “event” happens (i.e., the γ increases to a value that makes correlations become antiferromagnetic relative to before).

We sampled β linearly between β_min_ and β_max_, given that β_min_ = β(γ_min_) and β_max_ = β(γ_max_). We then inverted the relationship between γ and β to calculate the γ sample,

$$\gamma\left( \beta\right)=\frac{\sum_{{}_{\left( i,j \right)\in E^{-}\left( \beta\right)}} A_{ij}+\beta\left( \sum_{{}_{\left( i,j \right)\in E^{-}\left( \beta\right)}} A_{ij}- \sum_{{}_{\left( i,j \right)\in E^{-}\left( \beta\right)}} A_{ij} \right)}{\sum_{{}_{\left( i,j \right)\in E^{-}\left( \beta\right)}} P_{ij}+\beta\left( \sum_{{}_{\left( i,j \right)\in E^{-}\left( \beta\right)}} P_{ij}- \sum_{{}_{\left( i,j \right)\in E^{-}\left( \beta\right)}} P_{ij} \right)}$$

(5)

*Consensus Clustering –* Community detection algorithms based on modularity maximization suffer from degeneracy, that is, multiple near-optimal results. When applying these algorithms in a network repeatedly, outputs with a degree of difference are expected. Consensus clustering is a recursive procedure that uses this inherent variation to define the degree of agreement of these near-optimal solutions and work with it in order to reach a more meaningful network partition (46,47).

The basis of consensus clustering implementation is the co-classification matrix C_ij_, an *i x j matrix* where entries are the proportion each node pair is assigned to a same community t, given a set of partitions g,

$$C_{ij}\left( g \right)=\frac{1}{g}\sum\delta\left( g_{i}\left( t \right),g_{j}\left( t \right) \right).$$

(6)

We computed our co-classification matrices based on 1000 partitions using the γ range described above. In the co-classification matrix, values of 1 or 0 reflect the consensus that a given node pair always or never belong in the same community, respectively. The consensus clustering treats the co-classification matrix as a new network to find a consensus for all other values. One approach to find consensus is to apply a threshold to the weights and all values below it to zero (48). Another approach is to build a null model based on the co-classification distribution and incorporate it in the calculation of P in the modularity quality function (44,49). Here, we adopted this procedure as proposed by Jeub et al. (2018), which uses the modularity quality function to perform a hypothesis test

$$Q_{C}\left( \vec{g},\alpha\right)=\sum\left( C_{ij}-P_{ij}^{null}\left( \alpha\right) \right)\delta\left( g_{i},g_{j} \right),$$

(7)

where α is the significance level, and P_ij_^null^(α) is the highest value between the expected probability of *i* belonging to the same community of *j* and vice-versa (see *Null Model* below). In this hypothesis test, pairs of nodes that co-occur significantly less than can be explained by the null matrix P_ij_^null^ contribute negatively to the equation in (7) and are separated (see Jeub et al., 2018 for more details).

In order to obtain a consensus partition, we used an iterative procedure (48). The sequence of obtaining a set of partitions with the Louvain algorithm, calculating the co-classification matrix and identifying the links between communities with equation (7) was repeated until the co-classification matrix was binary, that is, until there was consensus as to which nodes co-occurred in the same community and which did not.

*Null Model –* We adopted a local permutation procedure to derive a co-classification matrix under the null model. A full description and rationale for the local permutation procedure can be found in the original publication (44). Briefly, after we obtained a set of 1000 partitions $\vec{g}$ from the signed Louvain-like algorithm (the same used to compute co-classification matrix), we fixed the community assignment of node *i* and permuted that of node *j*, keeping the number of communities and their size fixed, according to

$$P_{ij}^{null}\left( t \right)=Pr\left[ g_{j}^{0}\left( t \right)=g_{i}\left( t \right)\vee g_{i}^{0}\left( t \right)=g_{i}\left( t \right) \right]=\frac{\left[ \vec{g}\left( t \right)=g_{i}\left( t \right) \right]-1}{n-1},$$

(8)

where $\left[ \vec{g}\left( t \right)=g_{i}\left( t \right) \right]$ is the number of times node *i* is assigned to community *t*.

After the permutation procedure, we estimated the distribution and confidence intervals of the co-classification matrix under the null model using a pseudo-random sampling approximation at a level of significance of 0.05. Since this procedure keeps node *i* fixed and permutes node *j,* it assumes that $P_{ij}^{null}\neq P_{ji}^{null}$, for which we adopted the higher probability.

*Hierarchical Consensus Clustering (HCC) –* The consensus clustering procedure so far described the detection of communities at a single level. To fully assess all the levels of meaningful clustering in the networks, we made use of a recursive strategy proposed by (44). Briefly, once we detected the community structure at a given level, we treated each community as a network and repeated the procedure until no new communities were found at the significance level. This procedure generated a hierarchical clustering tree with multiple levels of significance.

After running HCC in the networks, we calculated the adjusted mutual information (AMI) between the final partition obtained from HCC or the anatomical groups partition and each partition of the partition distribution obtained using the γ range (Figure S1). We also show y(β) distribution used to compute those partitions (Figure S1).

**Network metrics**

All metrics were signed and normalized variants as suggested elsewhere (42). Briefly, given a node *i*, its strength *s* is defined as the sum of all connection weights in *i*, the within-community strength *s_i_(g_i_)* is the strength of node *i* within its community *g_i_*, and diversity coefficient is the normalized Shannon entropy of node *i* given its community *g_i_*. We calculated the positive and negative contributions to normalized strength as

$s_{i}'^{\pm}=\frac{s_{i}^{\pm}}{n-1}$,

(9)

Where *n* is the sum of the number of connections. The positive and negative contributions to normalized diversity coefficient were calculated as

$$h_{i}^{\pm}=-\frac{1}{log\left( max\left( g \right) \right)}\sum_{g_{i}\in g} p_{i}^{\pm}\left( u \right)logp_{i}^{\pm}\left( u \right),$$

(10)

Where max(*g)* is the number of communities in the community partition *g*,$p\left( u \right)=\frac{s_{i}^{\pm}\left( g_{i} \right)}{s_{i}^{\pm}}$, where $s_{i}^{\pm}\left( g_{i} \right)$is the within-community strength of node *i*, given its community *g_i_*. These normalized metrics vary in the range of [0,1]. Finally, we normalized the metrics such that positive contributions have more weight than negative,

| $s_{i}^{*}=s_{i}'^{+-}$, $h_{i}^{*}=h_{i}^{+-}$ | (11) |
| --- | --- |

Both $s_{i}^{*}$and $h_{i}^{*}$have a [-1,1] range. Note that the normalizations shown above for strength were applied to within-community strength, which was the metric used here. A detailed rationale about the normalization can be found in the original proposal (43)**.** The normalization allows a proper comparison of our metrics between networks. We compared these metrics across the conditions in our study as described above.
