## Supplementary figures and images for "Alcohol dependence modifies brain networks activated during abstinence and reaccess: a c-fos-based analysis in mice"

### Figure S1

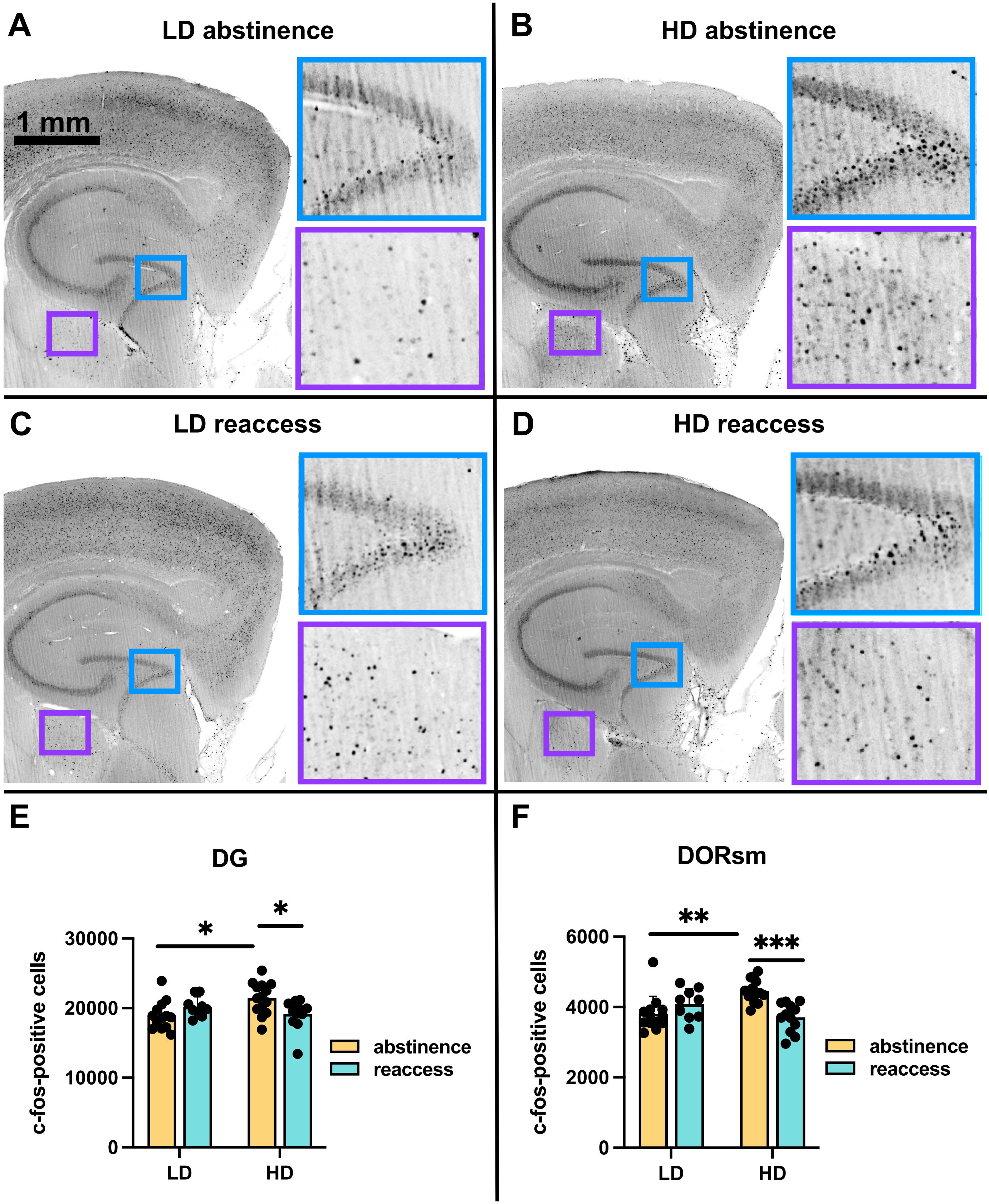

### Figure S2

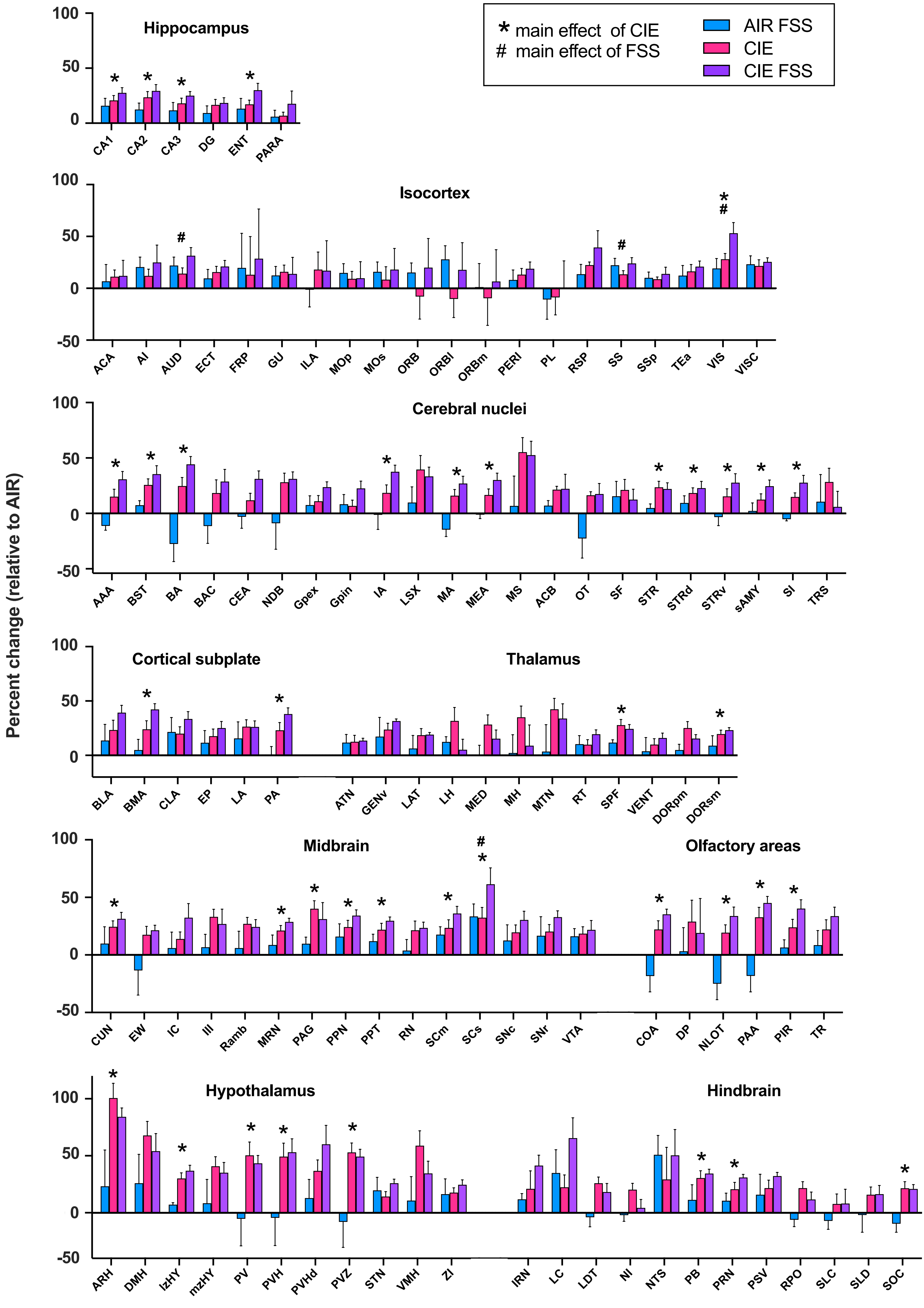

### Figure S3

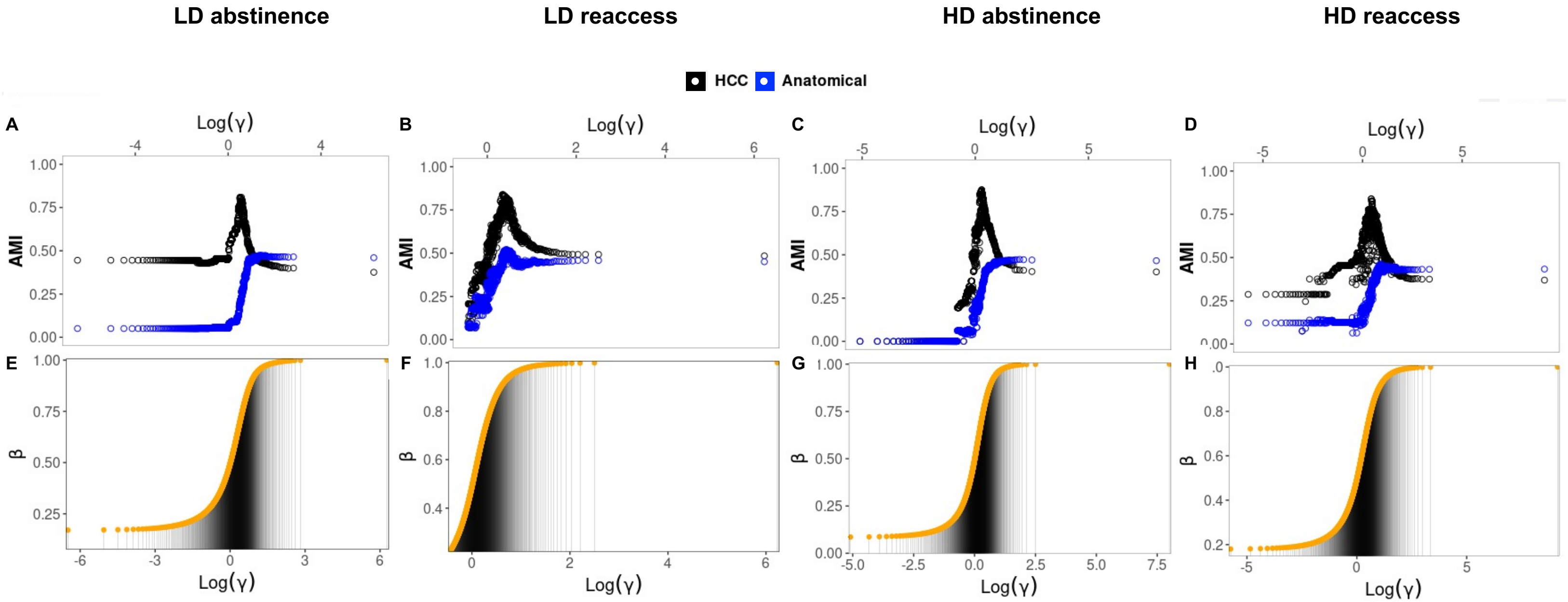
