## Supplementary material for "Alcohol dependence modifies brain networks activated during abstinence and reaccess: a c-fos-based analysis in mice": Table S1

**Supplemental Table 1: Global network metrics.**

| **Group** | **mean_FC** | **Anatomical**  **Modularity** | **HCC**  **Modularity** |
| --- | --- | --- | --- |
| **HD abstinence** | 0.2888 | 0.0341 | 0.1723 |
| **HD reaccess** | 0.1101 | 0.0829 | 0.2843 |
| **LD abstinence** | 0.1508 | 0.0496 | 0.3698 |
| **LD reaccess** | 0.2328 | 0.681 | 0.1518 |
