## Supplementary material for "Alcohol dependence modifies brain networks activated during abstinence and reaccess: a c-fos-based analysis in mice": Table S2

**Supplemental Table 2: Comparison of global level metrics among all four groups, two by two.**

| **Comparisons** | | **mean_FC** | | **Anatomical Modularity** | | **HCC Modularity** | |
| --- | --- | --- | --- | --- | --- | --- | --- |
|  |  | **Estimate** | **p value** | **Estimate** | **p value** | **Estimate** | **p value** |
| **HD abstinence** | **vs. HD reaccess** | 0.179 | 0.1815 | **0.049** | **0.0366** | **0.112** | **0.0502** |
| **HD abstinence** | **vs. LD abstinence** | 0.138 | 0.3002 | 0.015 | 0.5205 | 0.1975 | 0.1241 |
| **HD reaccess** | **vs. LD reaccess** | 0.123 | 0.4233 | 0.015 | 0.5555 | **0.1325** | **0.0116** |
| **LD abstinence** | **vs. LD reaccess** | 0.082 | 0.586 | 0.019 | 0.4539 | 0.218 | 0.166 |

Bold cells indicate significant differences at the significance level of α = 0.05
